# Neural Encoding of Auditory Features during Music Perception and Imagery: Insight into the Brain of a Piano Player *Auditory Features and Music Imagery*

**DOI:** 10.1101/106617

**Authors:** Stephanie Martin, Christian Mikutta, Matthew K. Leonard, Dylan Hungate, Stefan Koelsch, Edward F. Chang, José del R. Millán, Robert T. Knight, Brian N. Pasley

## Abstract

It remains unclear how the human cortex represents spectrotemporal sound features during auditory imagery, and how this representation compares to auditory perception. To assess this, we recorded electrocorticographic signals from an epileptic patient with proficient music ability in two conditions. First, the participant played two piano pieces on an electronic piano with the sound volume of the digital keyboard on. Second, the participant replayed the same piano pieces, but without auditory feedback, and the participant was asked to imagine hearing the music in his mind. In both conditions, the sound output of the keyboard was recorded, thus allowing precise time-locking between the neural activity and the spectrotemporal content of the music imagery. For both conditions, we built encoding models to predict high gamma neural activity (70-150Hz) from the spectrogram representation of the recorded sound. We found robust similarities between perception and imagery – in frequency and temporal tuning properties in auditory areas.

**Abbreviations:**

ECoG: electrocorticography
HG: high gamma
IFG: inferior frontal gyrus
MTG: middle temporal gyrus
Post-CG: post-central gyrus
Pre-CG: pre-central gyrus
SMG: supramarginal gyrus
STG: superior temporal gyrus
STRF: spectrotemporal receptive field

## Introduction

Auditory imagery is defined here as the mental representation of sound perception in the absence of external auditory stimulation. The experience of auditory imagery is common, such as when a song runs continually through someone’s mind. On an advanced level, professional musicians are able to imagine a piece of music by looking at the sheet music [1]. Behavioral studies have shown that structural and temporal properties of auditory features (see [2] for complete review), such as pitch [3], timbre [4,5], loudness [6] and rhythm [7] are preserved during auditory imagery. However, it is unclear how these auditory features are represented in the brain during imagery. Experimental investigation is difficult due to the lack of observable stimulus or behavioral markers during auditory imagery. Using a novel experimental paradigm to synchronize auditory imagery events to neural activity, we investigated the neural representation of spectrotemporal auditory features during auditory imagery in an epileptic patient with proficient music abilities.

Previous studies have identified anatomical regions active during auditory imagery [8], and how they compare to actual auditory perception. For instance, lesion [9] and brain imaging studies [4,10–14] have confirmed the involvement of bilateral temporal lobe regions during auditory imagery (see [15] for a review). Brain areas consistently activated with fMRI during auditory imagery include the secondary auditory cortex [10,12,16], the frontal cortex [16,17], the sylvian parietal temporal area [18], ventrolateral and dorsolateral cortices [19] and the supplementary motor area [17,20–24]. Anatomical regions active during auditory imagery have been compared to actual auditory perception to understand the interactions between externally and internally driven cortical processes. Several studies showed that auditory imagery has substantial, but not complete overlap in brain areas with music perception [8] – e.g. the secondary auditory cortex is consistently activated during music imagery and perception while the primary auditory areas appear to be activated solely during auditory perception [4,10,25,26].

These studies have helped to unravel anatomical brain areas involved in auditory perception and imagery, however, there is lack of evidence for the representation of specific acoustic features in the human cortex during auditory imagery. It remains a challenge to investigate neural processing during internal subjective experience like music imagery, due to the difficulty in time-locking brain activity to a measurable stimulus during auditory imagery. To address this issue, we recorded electrocorticographic neural signals (ECoG) of a proficient piano player in a novel task design that permitted robust marking of the spectrotemporal content of the intended music imagery to neural activity – thus allowing us to investigate specific auditory features during auditory imagery. In the first condition, the participant played an electronic piano with the sound output turned on. In this condition, the sound was played out loud through speakers at a comfortable sound volume that allowed auditory feedback (perception condition). In the second condition, the participant played the electronic piano with the speakers turned off, and instead imagined the corresponding music in his mind (imagery condition). In both conditions, the digitized sound output of the MIDI-compatible sound module was recorded. This provided a measurable record of the content and timing of the participant’s music imagery when the speakers of the keyboard were turned off and he did not hear the music. This task design allowed precise temporal alignment between the recorded neural activity and spectrogram representations of music perception and imagery – providing a unique opportunity to apply receptive field modeling techniques to quantitatively study neural encoding during auditory imagery.

A well-established role of the early auditory system is to decompose complex sounds into their component frequencies [27–29], giving rise to tonotopic maps in the auditory cortex (see [30] for a review). Auditory perception has been extensively studied in animal models and humans using spectrotemporal receptive field (STRFs) analysis [27,31–34], which identifies the time-frequency stimulus features encoded by a neuron or population of neurons. STRFs are consistently observed during auditory perception tasks, but the existence of STRFs during auditory imagery is unclear due to the experimental challenges associated with synchronizing neural activity and the imagined stimulus. To characterize and compare the spectrotemporal tuning properties during auditory imagery and perception, we fitted two encoding models on data collected from the perception and imagery conditions. In this case, encoding models describe the linear mapping between a given auditory stimulus representation and its corresponding brain response. For instance, encoding models have revealed the neural tuning properties of various speech features, such as acoustic, phonetic and semantic representations [33,35–38].

In this study, the neural representation of music perception and imagery was quantified by spectrotemporal receptive fields (STRFs) that predict high gamma (HG; 70-150Hz) neural activity. High gamma correlates with the spiking activity of the underlying neuronal ensemble [39–41] and reliably tracks speech and music features in auditory and motor cortex [33,42–45]. Results demonstrated the presence of robust spectrotemporal receptive fields during auditory imagery with extensive overlap in frequency tuning and cortical location compared to receptive fields measured during auditory perception. These results provide a quantitative characterization of the shared neural representation underlying auditory perception and the subjective experience of auditory imagery.

## Results

### High gamma neural encoding during auditory perception and imagery

In this study, ECoG recordings were obtained using subdural electrode arrays implanted in a patient undergoing neurosurgical procedures for epilepsy. The participant was a proficient piano player (age of start: 7, years of music education: 10, hours of training per week: 5). Prior studies have shown that music training is associated with improved auditory imagery ability, such as pitch and temporal acuity [46–48]. We took advantage of this rare clinical case to investigate the neural correlates of music perception and imagery. In the first task, the participant played two music pieces on an electronic piano with the speakers of the digital keyboard turned on (perception condition; Fig 1A). In the second task, the participant played the two same piano pieces, but the volume of the speaker system was turned off to establish a silent room. The participant was asked to imagine hearing the corresponding music in his mind as he played the piano (imagery condition; Fig 1B). In both conditions, the sound produced by the MIDI-compatible sound module was digitally recorded in synchrony with the ECoG signal. The recorded sound allowed synchronizing the auditory spectrotemporal patterns of the imagined music and the neural activity when the speaker system was turned off and no audible sounds were recorded in the room.

**Fig 1.**
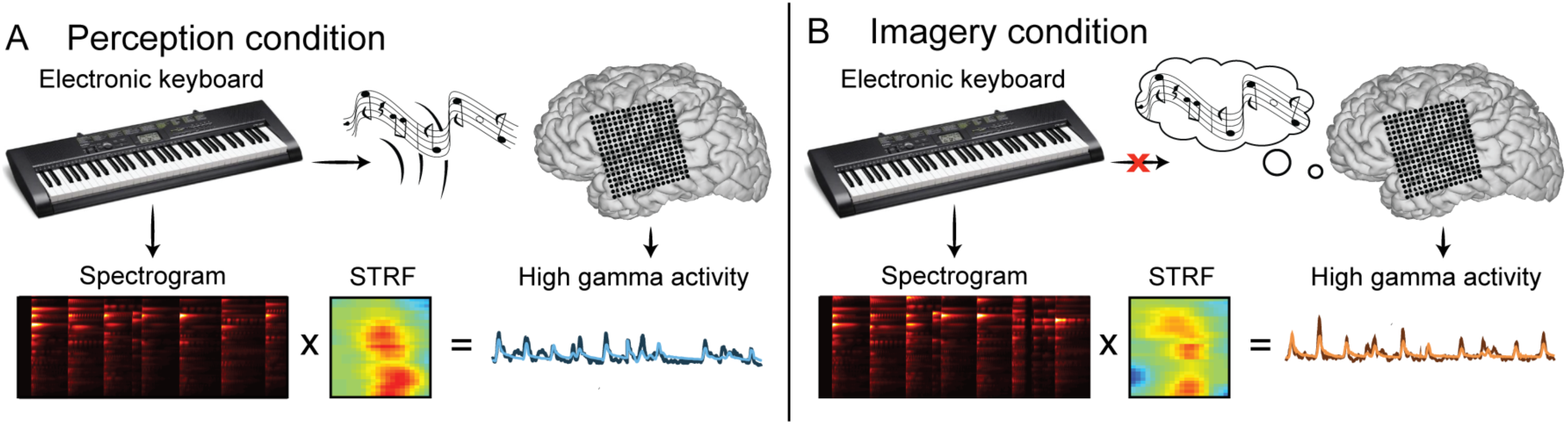
**Experimental task design.** (A) The participant played an electronic piano with the sound of the digital keyboard turned on (perception condition). (B) In the second condition, the participant played the piano with the sound turned off and instead imagined the corresponding music in his mind (imagery condition). In both conditions, the digitized sound output of the MIDI-compatible sound module was recorded in synchrony with the neural signals (even when the participant did not hear any sound in the imagery condition). The models take as input a spectrogram consisting of time-varying spectral power across a range of acoustic frequencies (200– 7,000 Hz, bottom left) and output time-varying neural signals. To assess the encoding accuracy, the predicted neural signal (light lines) is compared to the original neural signal (dark lines).

The participant performed two famous music pieces: Chopin’s Prelude in C minor, Op. 28, No.20 and Bach’s Prelude in C major BWV 846. Example auditory spectrograms from Chopin’s Prelude determined through the participant's key presses with the electronic piano are shown in Fig 2B for both perception and imagery conditions. To evaluate how consistently the participant performed across perception and imagery tasks, we computed the realigned Euclidean distance [49] between the spectrograms of the same music pieces played across conditions (within-stimulus distance). We compared the within-stimulus distance with the realigned Euclidean distance between the spectrograms of the different musical pieces (between-stimulus distance). The realigned Euclidean distance was 251.6% larger for the between-stimulus distance compared to the within-stimulus distance (p<10^-3^; randomization test), suggesting that the spectrograms of same musical pieces played across conditions were more similar than the spectrograms of different musical pieces. This indicates that the participant performed the task with relative consistency and specificity across the two conditions. To compare spectrotemporal auditory representations during music perception and music imagery tasks, we fit separate encoding models in each condition. We used these models to quantify specific anatomical and neural tuning differences between auditory perception and imagery.

**Fig 2.**
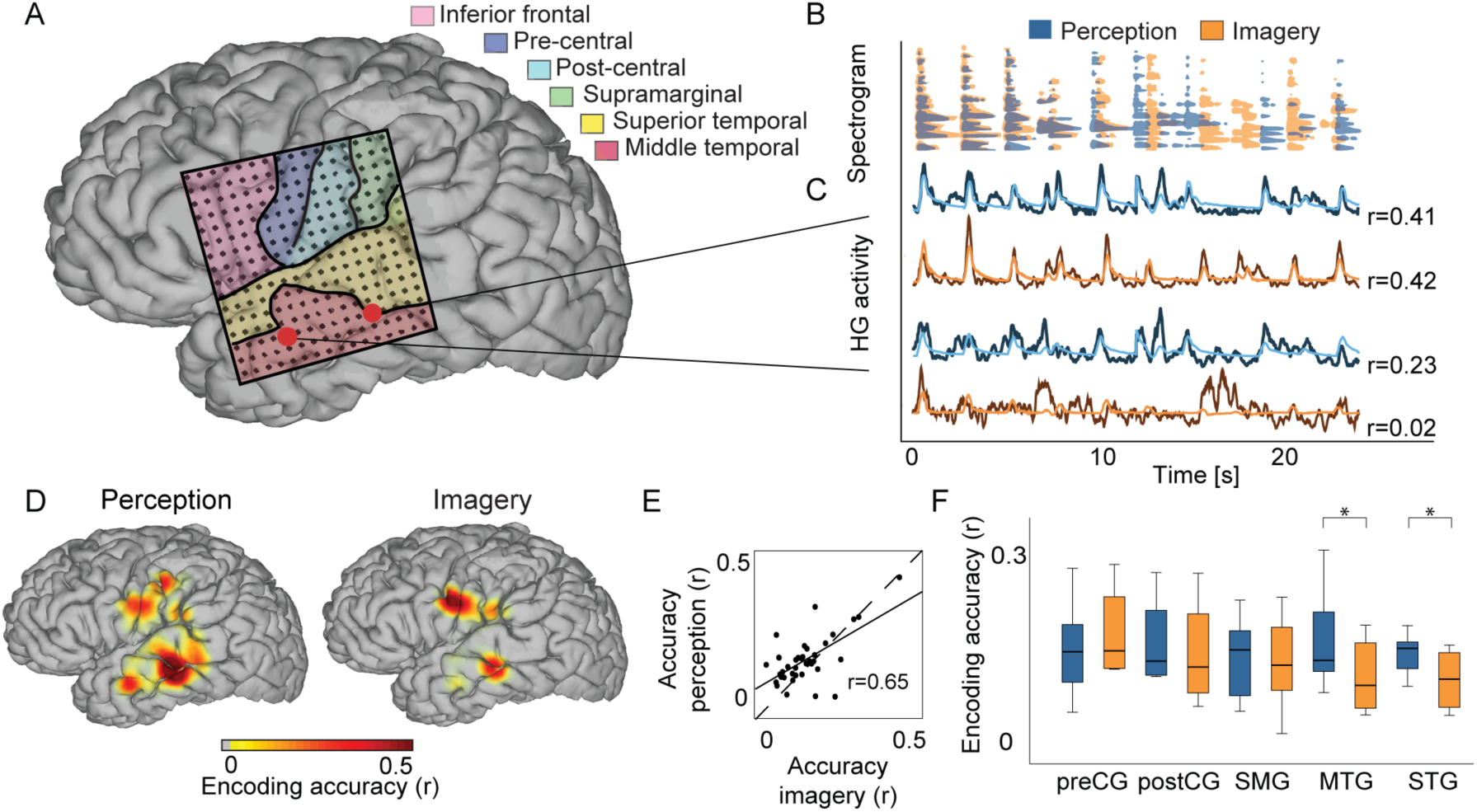
**Encoding accuracy.** (A) Electrode location overlaid on cortical surface reconstruction of the participant’s cerebrum. (B) Overlay of the spectrogram contours for the perception (blue) and imagery (orange) condition (10% of maximum energy from the spectrograms). (C) Actual and predicted high gamma band power (70–150 Hz) induced by the music perception and imagery segment in (B). Top electrodes have very similar predictive power, whereas bottom electrodes are very different for perception and imagery. Recordings are from two different STG sites, highlighted in pink in (A). (D) Encoding accuracy is plotted on the cortical surface reconstruction of the participant’s cerebrum (map thresholded at p<0.05; FDR correction). (E) Encoding accuracy of significant electrodes of the perception model as a function the imagery model. Electrode-specific encoding accuracy is correlated between both perception and imagery models (r=0.65; p<10^-4^; randomization test). (F) Encoding accuracy as a function of anatomic location (pre-central gyrus (pre-CG), post-central gyrus (post-CG), supramarginal gyrus (SMG), medial temporal gyrus (MTG) and superior temporal gyrus (STG)).

For both perception and imagery conditions, the observed and predicted high gamma neural responses are illustrated for two individual electrodes in the temporal lobe, respectively (Fig 2C), together with the corresponding music spectrum (Fig 2B). The predicted neural response for the electrode shown in the upper panel of Fig 2B was significantly correlated with its corresponding measured neural response in both perception (r=0.41; p<10^-7^; one-sample *z*-test; FDR correction) and imagery (r=0.42; p<10^-4^; one-sample *z*-test; FDR correction) conditions. The predicted neural response for the lower panel electrode was correlated with the actual neural response only in the perception condition (r=0.23; p<0.005; one-sample *z*-test; FDR correction) but not in the imagery condition (r=-0.02; p>0.5; one-sample *z*-test; FDR correction). The difference between both conditions was significant for the electrode in the lower panel (p<0.05; two-sample t-test), but not in the upper panel (p>0.5; two-sample t-test). This suggests that there is a strong continuous relationship between time-varying imagined sound features and STG neural activity, but that this relationship is dependent on cortical location.

To further investigate anatomical similarities and differences between the perception and imagery conditions, we plotted the anatomical layout of prediction accuracy of individual electrodes. In both conditions, results showed that sites with the highest prediction in both conditions were located in the superior and middle temporal gyrus, pre-and post-central gyrus, and supramarginal gyrus (Fig 2D; heat map thresholded to p<0.05; one-sample *z*-test; FDR correction), consistent with previous ECoG results [33]. Among the 256 electrodes recorded, 210 were fitted in the encoding model, while the remaining 46 electrodes were removed due to excessive noise. Within the fitted electrodes, while 35 and 15 electrodes were significant in the perception and imagery condition, respectively (p<0.05; one-sample *z*-test; FDR correction), of which 9 electrodes were significant in both conditions. Anatomic locations of the electrodes with significant encoding accuracy are depicted in Fig 3 and Fig S1. To compare the encoding accuracy across conditions, we performed additional analysis on the electrodes that had significant encoding accuracy in at least one condition (41 electrodes; unless otherwise stated). Prediction accuracy of individual electrodes was correlated between perception and imagery (Fig 2E; 41 electrodes; r=0.65; p<10^-4^; randomization test). Because both perception and imagery models are based on the same auditory stimulus representation, the correlated prediction accuracy provides strong evidence for a shared neural representation of sound based on spectrotemporal features.

To assess how brain areas encoding auditory features varied across experimental conditions, we analyzed the significant electrodes in the gyri highlighted in Fig 2A (pre-central gyrus (pre-CG), post-central gyrus (post-CG), supramarginal gyrus (SMG), medial temporal gyrus (MTG) and superior temporal gyrus (STG)) using Wilcoxon signed-rank test (p>0.05; one-sample Kolmogorov-Smirnov test; Fig 2F). Results showed that the encoding accuracy in the MTG and STG was higher for the perception (MTG: *M* = 0.16, STG: *M* = *0.13)* than for the imagery (MTG: *M* = 0.11, STG: *M*= 0.08; p<0.05; Wilcoxon signed-rank test; Bonferroni correction). The encoding accuracy in the pre-CG, post-CG and SMG was not different between the perception (pre-CG *M*=0.17; post-CG *M*=0.15; SMG *M*=0.12; p>0.5; Wilcoxon signed-rank test; Bonferroni correction) and imagery (pre-CG *M*=0.14; post-CG *M*=0.13; SMG *M*=0.12; p>0.5; Wilcoxon signed-rank test; Bonferroni correction) conditions. The significant improvement of the perception vs. imagery model was thus specific to the temporal lobe, which may reflect underlying differences in spectrotemporal encoding mechanisms, or alternatively, a greater sensitivity to discrepancies between the actual content of imagery and the recorded sound stimulus used in the model.

### Spectrotemporal tuning during auditory perception and imagery

Auditory imagery and perception are distinct subjective experiences yet both are characterized by a sense of sound. How does the auditory system encode sound during music perception and imagery? Examples of standard STRFs are shown in Fig 3 for temporal electrodes (Fig S2 for all the STRFs). These STRFs highlight neural stimulus preferences as shown by the excitatory (warm color) and inhibitory (cold color) subregions. Fig 4A shows the latency (s) for the perception and imagery conditions, defined as the temporal coordinates of the maximum deviation in the STRF. The peak latency was significantly correlated between both conditions (r=0.43; p<0.005; randomization test), suggesting that both perception and imagery are simultaneously active for most of their response durations. Then, we analyzed frequency tuning curves estimated from the STRFs (see Materials and Methods for details). Examples of tuning curves for both perception and imagery encoding models are shown for the electrodes indicated by the black outline in the anatomic brain (Fig 4B). Across conditions, the majority of individual electrodes exhibited a complex frequency tuning profile. For each electrode, similarities between the tuning curves in the perception and imagery models were quantified using Pearson’s correlation coefficient. The anatomical distribution of tuning curve similarity is plotted in Fig 4B, with the correlation at individual sites ranging between r=-0.3-0.6. The effect of anatomical location (pre-CG, post-CG, SMG, MTG and STG) on tuning curve similarity was not significant (*Chi-square*=3.59; p>0.1; Kruskal-Wallis Test). Similarities in tuning curve shape between auditory imagery and perception suggest a shared auditory representation, but there is no evidence that similarities depended on gyral area.

**Fig 3.**
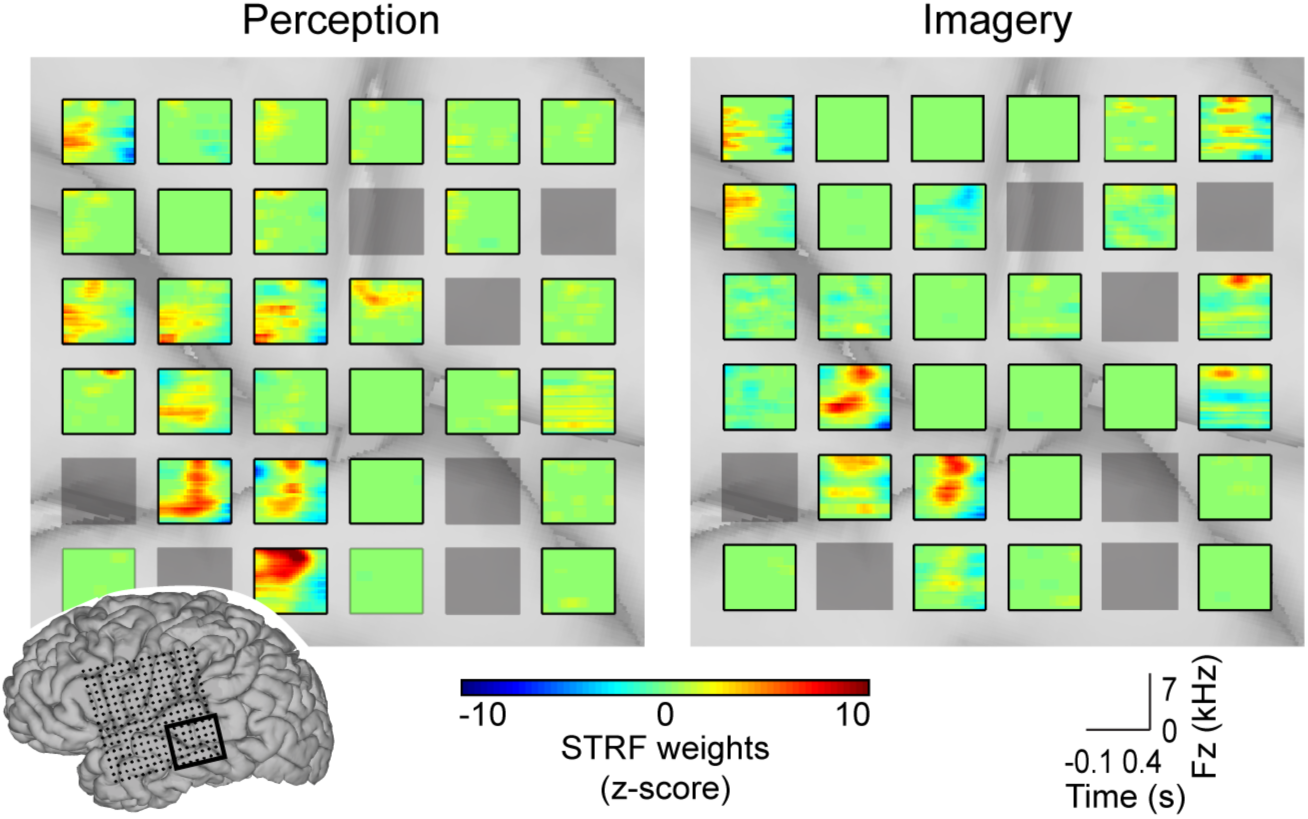
**Spectrotemporal receptive fields (STRFs)**. Examples of standard STRFs for the perception (left panel) and imagery (right panel) models (warm colors indicate where the neuronal ensemble is excited, cold colors indicate where the neuronal ensemble is inhibited). On the lower left corner, Electrode location overlaid on cortical surface reconstructions of the participant’s cerebrum. Electrodes whose STRFs are shown are outlined in black. Grey electrodes were removed from the analysis due to excessive noise (see Materials and Methods).

**Fig 4.**
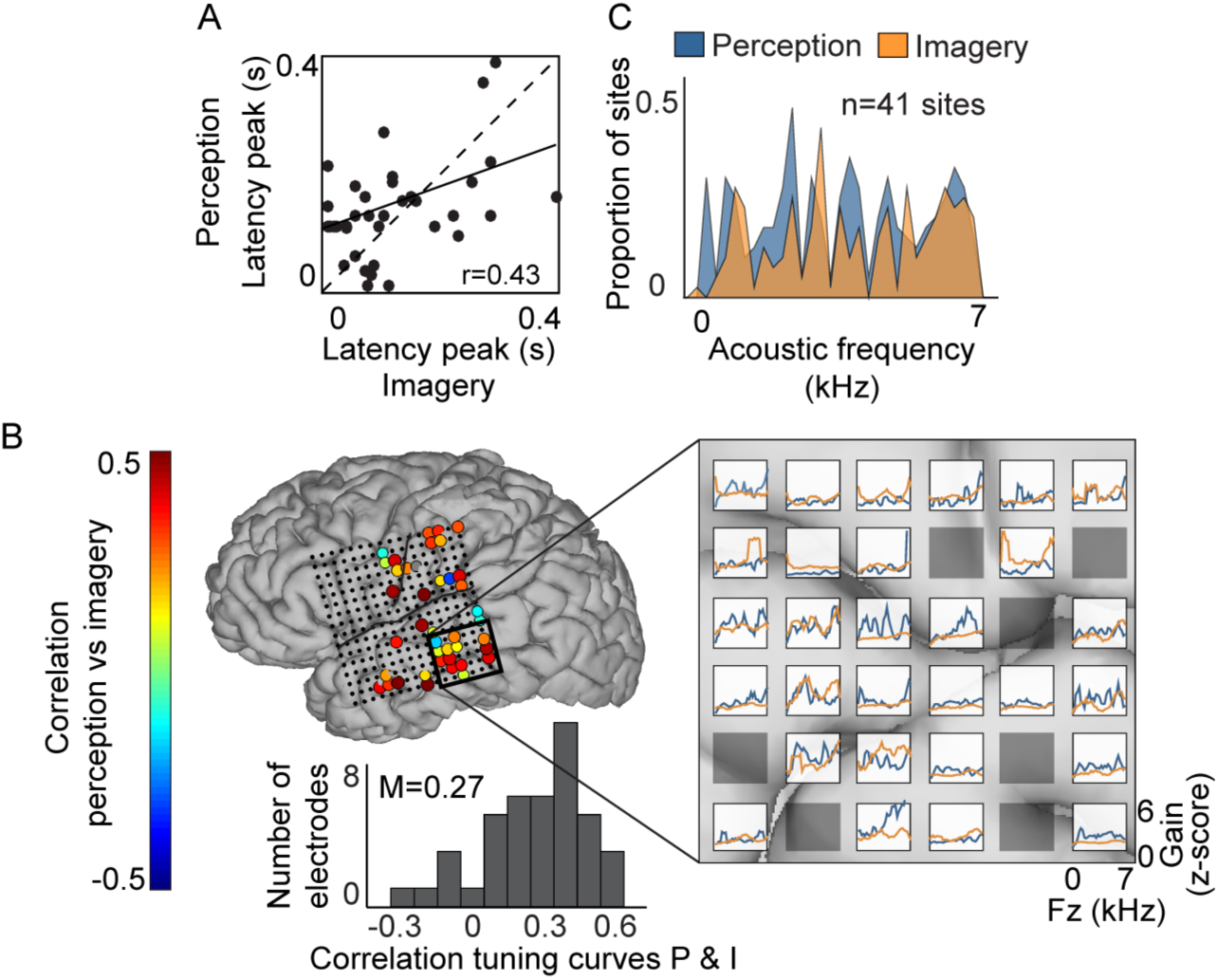
**Auditory tuning**. (A) Latency peaks – estimated from the STRFs – were significantly correlated between perception and imagery conditions (r=0.43; p<0.005; randomization test). (B) Examples of tuning curves for both perception and imagery encoding models defined as the average gain of the STRFs as a function of acoustic frequency. Black outline in the anatomic brain indicate electrode location. for the electrodes indicated by the black outline in the left panel – Correlation coefficients between the perception and imagery conditions are plotted for significant electrodes on the cortical surface reconstruction of the participant’s cerebrum. Grey electrodes were removed from the analysis due to excessive noise. Bottom panel is a histogram of electrode correlation coefficients between the perception and imagery tuning. (C) Proportion of predictive electrode sites (N=41) with peak tuning at each frequency. Tuning peaks were identified as significant parameters in the acoustic frequency tuning curves (z>3.1; p<0.001) – separated by more than one third an octave.

Different electrodes are sensitive to different acoustic frequencies important for music processing. We next quantitatively assessed how frequency tuning of predictive electrodes (N=41) varied during the two conditions. First, to evaluate how the acoustic spectrum was covered at the population level, we quantified the proportion of significant electrodes with a tuning peak at each acoustic frequency (Fig 4C). Tuning peaks were identified as significant parameters in the acoustic frequency tuning curves (z>3.1; p<0.001; separated by more than one third an octave). The proportion of electrodes with tuning peaks was significantly larger for the perception (mean = 0.19) than for the imagery (mean = 0.14) condition (Fig 4C; p<0.05; Wilcoxon signed-rank test). Despite this, both conditions exhibited reliable frequency selectivity, as nearly the full range of the acoustic frequency spectrum was encoded. The fraction of acoustic frequency covered with peaks by predictive electrodes was 0.91 for the perception and 0.89 for the imagery.

### Reconstruction of auditory features during music perception and imagery

To evaluate the ability to identify piano keys from the brain activity, we reconstructed the same auditory spectrogram representation used in the encoding models. Results showed that the overall reconstruction accuracy was higher than zero in both conditions (Fig 5A; p < 0.001; randomization test), but did not differ between conditions (p > 0.05; two-sample t-test). As a function of acoustic frequency, mean accuracy ranged from r=0–0.45 (Fig 5B).

**Fig 5.**
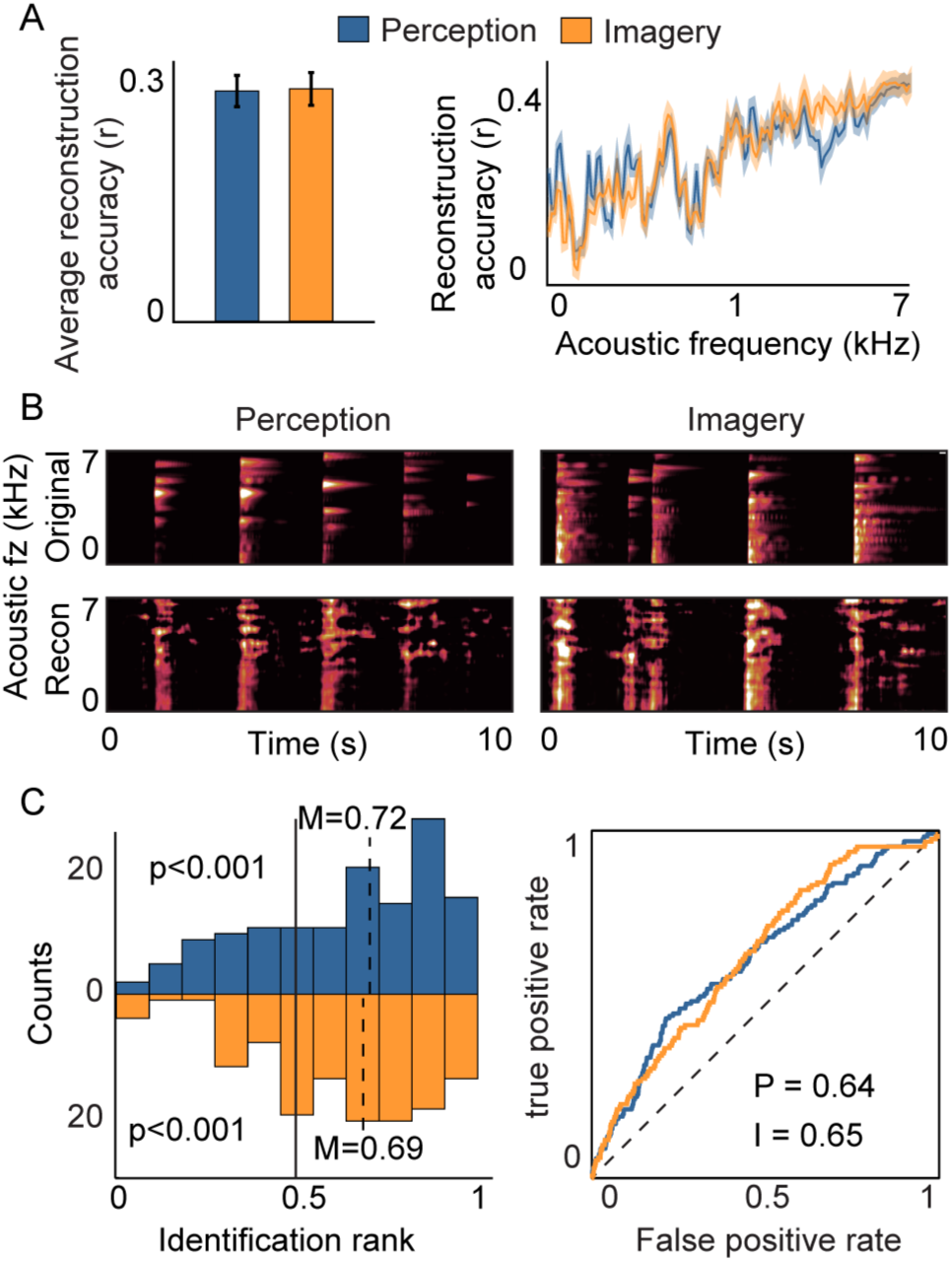
**Reconstruction accuracy**. (A) Right panel: Overall reconstruction accuracy of the spectrogram representation for both perception (blue) and imagery (orange) conditions. Error bars denote resampling SEM. Left panel: Reconstruction accuracy as a function of acoustic frequency. Shaded region denotes SEM over the resamples. (B) Examples of original and reconstructed segments for the perception (left) and the imagery (right) model. (C) Distribution of identification rank for all reconstructed spectrogram notes. Median identification rank is 0.65 and 0.63 for the perception and imagery decoding model, respectively, which is significantly higher than 0.50 chance level (p<0.001; randomization test). Left panel: Receiver operating characteristic (ROC) plot of identification performance for the perception (blue curve) and imagery (orange curve) model. Diagonal black line indicates no predictive power.

We further assessed reconstruction accuracy by evaluating the ability to identify isolated piano notes from the test set auditory spectrogram reconstructions. Examples of original and reconstructed segments are depicted in Fig 5B for the perception (left) and imagery model (right). For the identification, we extracted 0.5-second segments at piano note onsets from the original and reconstructed auditory spectrogram. Onsets of the notes were defined with the MIRtoolbox [50]. Then, we computed the correlation coefficient between a target reconstructed spectrogram and original spectrograms in the candidate set. Finally, we sorted the coefficients and computed the identification rank as the percentile rank of the correct spectrogram. This metric reflects how well the target reconstruction matched the correct original spectrogram out of all candidate. Results showed that the median identification rank of individual piano notes was significantly higher than chance for both conditions (Fig 5C; median identification rank perception = 0.72 and imagery = 0.69; p< 0.001; randomization test). Similarly, the area under the curve (AUC) of identification performance for the perception (blue curve) and imagery (orange curve) model was well above chance level (diagonal black dashed line indicates no predictive power; p<0.001; randomization test).

### Cross-condition analysis

Another way to evaluate the overlapping degree between both perception and imagery conditions is to apply the decoding model built in the perception condition to imagery neural data, and vice-versa. This approach is based on the hypothesis that both tasks share neural mechanisms and is useful when one of the models cannot be built directly, because of the lack of observable measures. This technique has been successfully applied to various fields, such as vision [51–53], and speech [54]. When the model was trained on the perception condition and tested on the imagined condition (r=0.28), decoding performances improved by 50% compared to when the perception model was applied to imagined data (r=0.19) (Fig 6). This highlight the importance of having a model that is specific to each condition.

**Fig 6.**
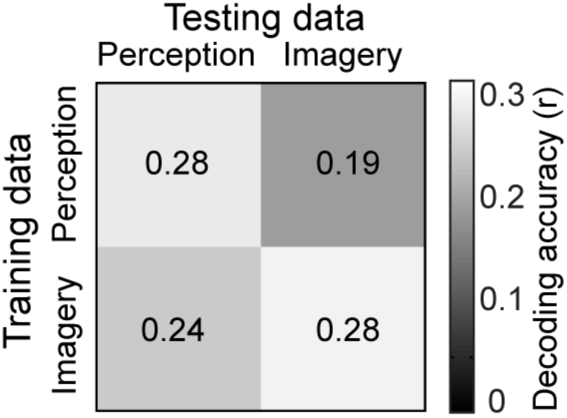
**Cross-condition analysis**. Reconstruction accuracy when the decoding model was built on the perception condition and applied to the imagery neural data and vis-versa. Decoding performances improved by 50% when the model was trained on the perception condition and tested on the imagined condition (r=0.28), compared to when the perception model was applied to imagined data (r=0.19).

### Control analysis for sensorimotor confounds

In this study, the participant played piano in two different conditions (music perception and imagery). In addition to the auditory percept, arm-, hand-and finger-movements related to the active piano task could have presented potential confounds to the decoding process. We controlled for possible motor confounds in three different ways. First, the electrode grid did not cover hand or finger sensorimotor brain area (Fig 7A). This reduces the likelihood that the decoding model involved hand-related motor confounds. Second, examples of STRFs for two neighboring electrodes show different weight patterns for both conditions (Fig 7B). For instance, the weights of the electrode depicted in the left example of Fig 7B are correlated between both conditions (r=0.48), whereas the weights are not correlated in the right example (r=0.04). Differences across conditions cannot be explained by motor confounds, because finger movement was similar in both tasks. Third, brain areas that significantly encoded music perception and imagery overlapped with auditory sensory areas (Fig S1 and Fig S3), as revealed by the encoding accuracy and STRFs during passive listening to TIMIT sentences (no movements). These findings provide evidence against motor confounds, and suggest that the brain responses were induced by auditory percepts rather than motor movements associated with pressing piano keys. Finally, we built two additional decoding models, using 1) only temporal lobe electrodes and 2) only auditory-responsive electrodes (Fig 7C; see Materials and Methods for details). Both models showed significant reconstruction accuracy (p<0.001; randomization test) and median identification rate (p<0.001; randomization test). This suggests that even if we removed all electrodes that are potentially not related to auditory processes, the decoding model still performs well.

**Fig 7.**
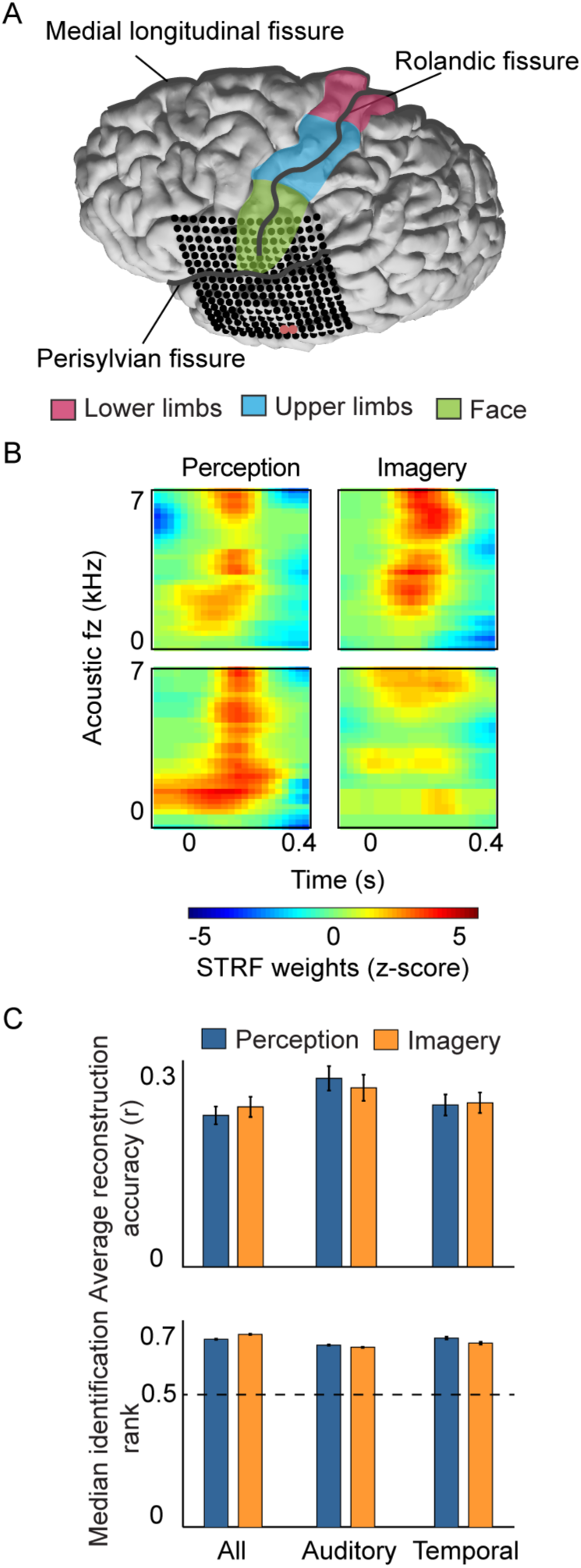
**Control analysis for motor confound**. (A) Electrode location overlaid on cortical surface reconstructions of the participant’s cerebrum. View from the top of the brain shows that hand motor areas not recorded with the grid. Brain areas associated with face, upper and lower limbs are depicted in green, blue and pink, respectively. (B) Different STRFs for two neighboring electrodes highlighted in (A) for both perception and imagery encoding models. (C) Overall reconstruction accuracy (left panel) and median identification rank (right panel) when using all electrodes, only temporal electrodes, and auditory electrodes (see Materials and methods for details).

## Discussion

Music imagery studies are daunting due to the subjective nature and absence of verifiable and observable measures. This task design allowed precise time-locking between the recorded neural activity and spectrotemporal features of music imagery, and provided a unique opportunity to quantitatively study neural encoding during auditory imagery, and compare tuning properties with auditory perception. We provide the first evidence of spectrotemporal receptive field and auditory features encoding in the brain during music imagery, and provided comparative measures with actual music perception encoding. We observed that neuronal ensembles were tuned to acoustic frequencies during imagined music, suggesting that spectral organization occurs in the absence of actual perceived sound. Supporting evidence has shown that restored speech – when a speech instance is replaced by noise, but the listener perceives a specific speech sound – is grounded in acoustic representations in the superior temporal gyrus [55,56]. In addition, results showed substantial, but not complete overlap in neural properties – i.e. spectral and temporal tuning properties, and brain areas – during music perception and imagery. These findings are in agreement with visual studies showing that visual imagery and perception involve many, but not all, overlapping brain areas [57,58]. We also showed that auditory features could be reconstructed from neural activity of the imagined music. Because both perception and imagery models are based on the same auditory stimulus representation, the correlated prediction accuracy provides strong evidence for a shared neural representation of sound based on spectrotemporal features. This confirms that the brain encodes spectrotemporal properties of sounds – as previously shown by behavioral and brain lesion studies (see [2] for a review).

Methodological issues in investigating imagery are numerous, including the lack of evidence that the desired mental task was operational. The current task design did not allow verifying how the mental task was performed, yet the behavioral index of keynote press on the piano indicated the precise time and frequency content of the intended imagined sound. In addition, we recorded a skilled piano player, and it has been suggested that participants with musical training exhibited better pitch and temporal acuity during auditory imagery than did participants with little or no musical training [47,59]. Furthermore, tonotopic maps located in the STG are enlarged within trained musicians [60]. Thus, having a trained piano-player suggests improved auditory imagery ability (see also [7,9,14], and reduced issues related to spectral and temporal errors.

Finally, the electrode grid was located on the left hemisphere of the participant. This raises the question of lateralization in the brain response to music perception and imagery. Studies have shown the importance of both hemispheres for auditory perception and imagination [4,9,10,12–14], although brain patterns tend to shift toward the right hemisphere (see [15] for a review). In our task, the grid was located on the left hemisphere, and allowed significant encoding and decoding accuracy within high gamma frequency ranges. This is consistent with the notion that music auditory processes are also evident in the left hemisphere.

## Materials and Methods

### Participant and data acquisition

Electrocorticographic (ECoG) recording was obtained using subdural electrode arrays implanted in one patient undergoing neurosurgical treatment for refractory epilepsy. The participant gave his written informed consent prior to surgery and experimental testing. The experimental protocol was approved by the University of California, San Francisco and Berkeley Institutional Review Boards and Committees on Human Research. Inter-electrode spacing (center-to-center) was 4 mm. Grid placement (Fig 2A) was defined solely based on clinical requirements. Localization and co-registration of electrodes was performed using the structural MRI. Multi-electrode ECoG data were amplified and digitally recorded with sampling rate of 3,052 Hz. ECoG signals were re-referenced to a common average after removal of electrodes with epileptic artifacts or excessive noise (including broadband electromagnetic noise from hospital equipment or poor contact with the cortical surface). In addition to the ECoG signals, the audio output of the piano was recorded along with the multi-electrode ECoG data.

### Experimental paradigm

The recording session included two conditions. In the first condition, the participant played on an electronic piano with the sound turned on. That is, the music was played out loud through the speakers of the digital keyboard in the hospital room (volume at comfortable and natural sound level; Fig 1A; perception condition). In the second condition, the participant played on the piano with the speakers turned off and instead imagined hearing the corresponding music in his mind (Fig 1B; imagery condition). Fig 1B illustrates that in both conditions, the digitized sound output of the MIDI sound module was recorded in synchrony with the ECoG data (even when the speakers were turned off and the participant did not hear the music). The two music pieces were Chopin’s Prelude in C minor Op. 28 no. 20 and Bach’s Prelude in C major (BWV 846), respectively.

### Feature extraction

We extracted the ECoG signal in the high gamma frequency band from eight bandpass filters (hamming window non-causal filter of order 20, logarithmically increasing center frequencies (70–150 Hz) and semi-logarithmically increasing bandwidths), and extracted the envelope using the Hilbert transform. Prior to model fitting, the power was averaged across these eight bands, down-sampled to 100 Hz and z-scored.

### Auditory spectrogram representation

The auditory spectrogram representation was a time-varying representation of the amplitude envelope at acoustic frequencies logarithmically spaced between 200-7,000 Hz. This representation was calculated by affine wavelet transforms of the sound waveform using auditory filter banks that mimics neural processing in the human auditory periphery [31]. To compute these acoustic representations, we used the NSL MATLAB toolbox (http://www.isr.umd.edu/Labs/NSL/Software.htm).

### Encoding model

The neural encoding model, based on the spectrotemporal receptive field (STRF) [34] describes the linear mapping between the music stimulus and the high gamma neural response at individual electrodes. The encoding model was estimated as follows:

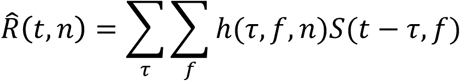

where 
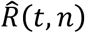
 is the predicted high gamma neural activity at time *t* and electrode *n*, *S(t-τ, f)* is the spectrogram representation at time *(t-τ)* and acoustic frequency *f*. Finally, *h(τ, f, n)* is the linear transformation matrix that depends on the time lag *τ*, the frequency *f* and electrodes *n*. *h* represents the spectrotemporal receptive field of each electrode. The Neural tuning properties of a variety of stimulus parameters in different sensory systems have been assessed using STRFs [61]. We used Ridge regression to fit the encoding model [62], and a 10-fold cross-validation resampling procedure, with no overlap between training and test partitions within each resample. We performed grid search on the training set to define the penalty coefficient *α* and the learning rate *η*, using a nested loop cross-validation approach. We standardized the parameter estimates to yield the final model.

### Decoding model

The decoding model linearly mapped the neural activity to the music representation, as a weighted sum of activity at each electrode, as follows:

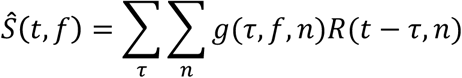

where 
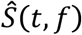
 is the predicted music representation at time *t* and frequency *f*. *R(t-τ, n)* is the HG neural response of electrode *n* at time *(t-τ)*, *τ* is the time lag ranging between −100ms and 400ms. Finally, *g(τ, f, n)* is the linear transformation matrix that depends on the time lag *τ*, frequency *f*, and electrode *n*. Both, neural response and music representation were synchronized, downsampled to 100 Hz, and standardized to zero mean and unit standard deviation prior to model fitting. To fit model parameters, we used gradient descent with early stopping regularization. We used a 10-folds cross-validation resampling scheme, and 20% of the training data were used as validation set to determine the early stopping criterion. Finally, model prediction accuracy was evaluated on the independent testing set, and the parameter estimates were standardized to yield the final model.

## Evaluation

Prediction accuracy was quantified using the correlation coefficient (Pearson’s r) between the predicted and actual HG signal using data from the independent test. Overall encoding accuracy was reported as the mean correlation over folds. The *z*-test was applied for all reported mean r values. Electrodes were defined as significant if the p-value was smaller than the significance threshold of *α*=0.05 (95th-percentile; FDR correction). To define auditory sensory areas, we built an encoding model on data recorded while the participant listened passively to speech sentences from the TIMIT corpus [63] during 10min. Electrodes with significant encoding accuracy are highlighted in Fig S1.

To further investigate the neural encoding of spectrotemporal acoustic features during music perception and music imagery, we analyzed all the electrodes that were at least significant in one condition (unless otherwise stated). Tuning curves were estimated from spectrotemporal receptive fields, by first setting all inhibitory weights to zero, then averaging across the time dimension and converting to standardized z-scores. Tuning peaks were identified as significant peak parameters in the acoustic frequency tuning curves (z>3.1; p<0.001) – separated by more than one third an octave.

Decoding accuracy was assessed by calculating the correlation coefficient (Pearson’s r) between the reconstructed and original music spectrogram representation using testing set data. Overall reconstruction accuracy was computed by averaging over acoustic frequencies and resamples, and standard error of the mean (SEM) was computed by taking the standard deviation across resamples. Finally, to further assess the reconstruction accuracy, we evaluated the ability to identify isolated piano notes from the test set auditory spectrogram reconstructions – using similar approach as in [33,54].

## Acknowledgments

This work was supported by the Zeno-Karl Schindler Foundation, NINDS Grant R3721135, NIH 5K99DC012804 and the Nielsen Corporation.

## Supporting Information

**Figure S1.**
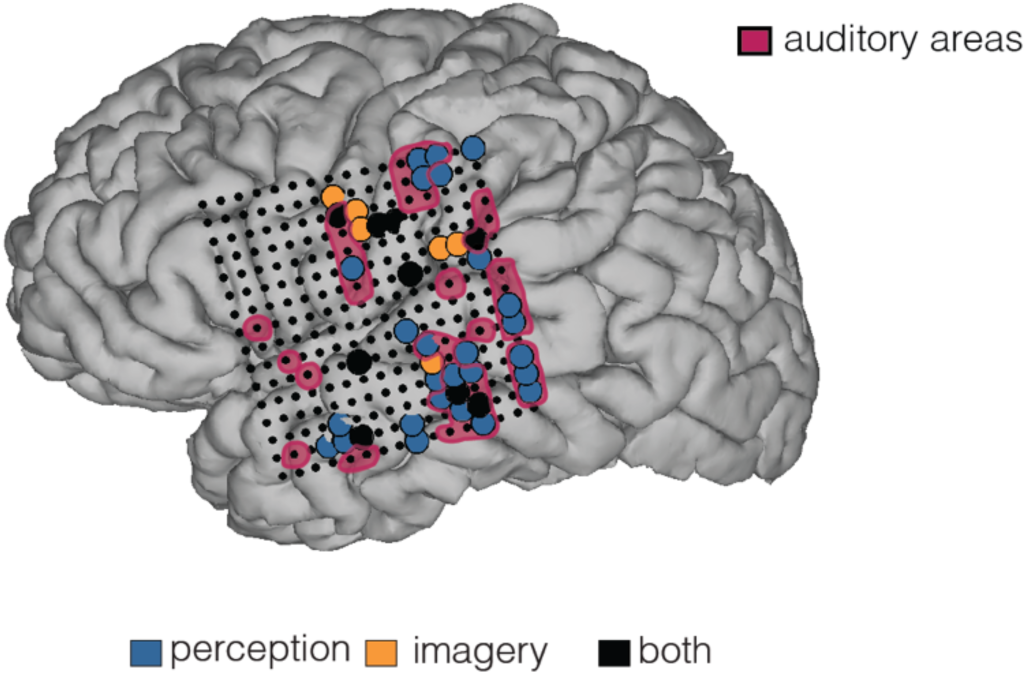
**Anatomical distribution of significant electrodes**. Electrodes with significant encoding accuracy overlaid on cortical surface reconstruction of the participant’s cerebrum. To define auditory sensory areas (pink), we built an encoding model on data recorded while participant listened passively to speech sentences from the TIMIT corpus [64]. Electrodes with significant encoding accuracy (p<0.05; FDA correction) are highlighted.

**Figure S2.**
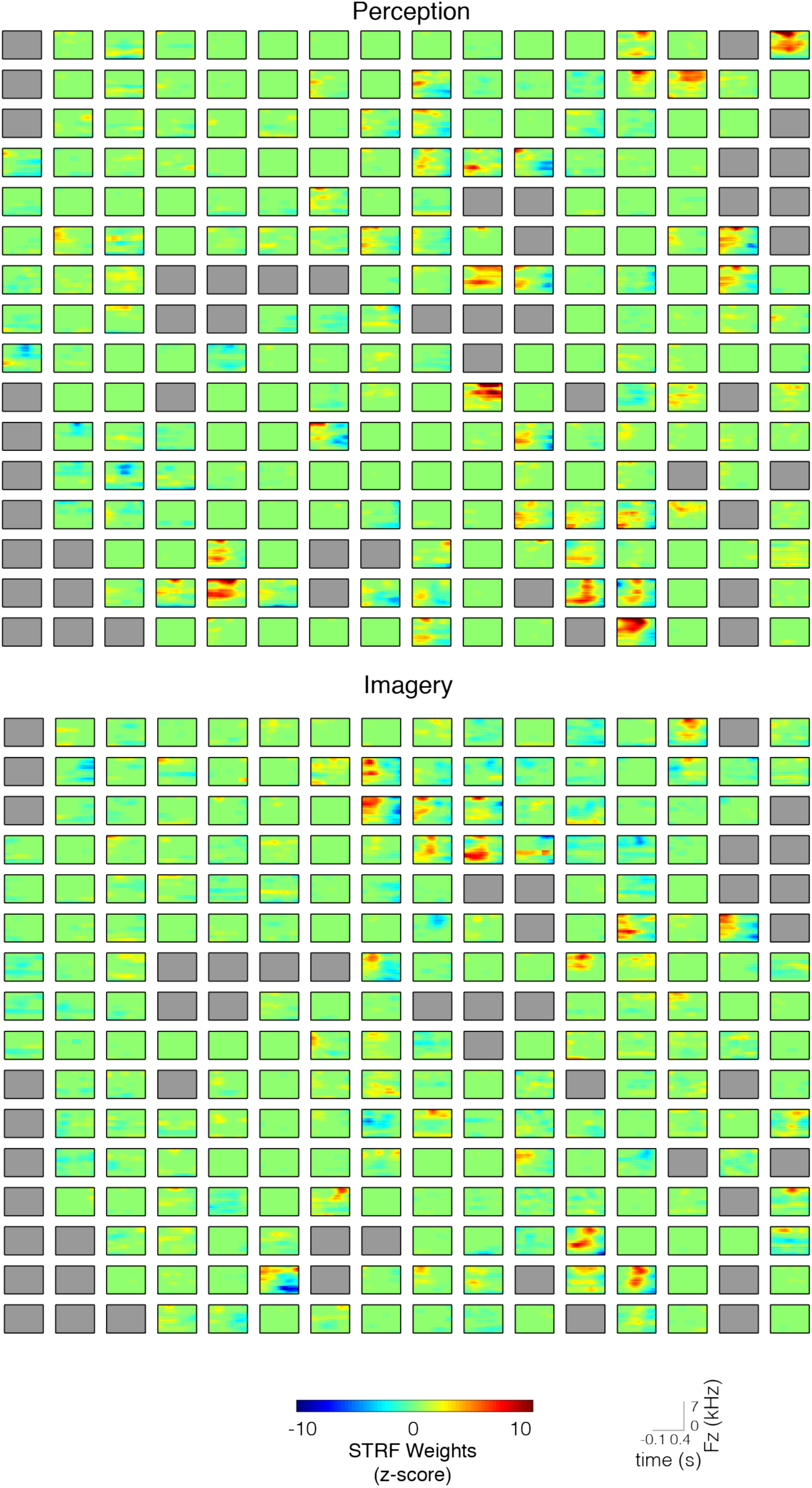
**Spectrotemporal receptive fields**. STRFs for the perception (top panel) and imagery (bottom panel) models. Grey electrodes were removed from the analysis due to excessive noise.

**Figure S3.**
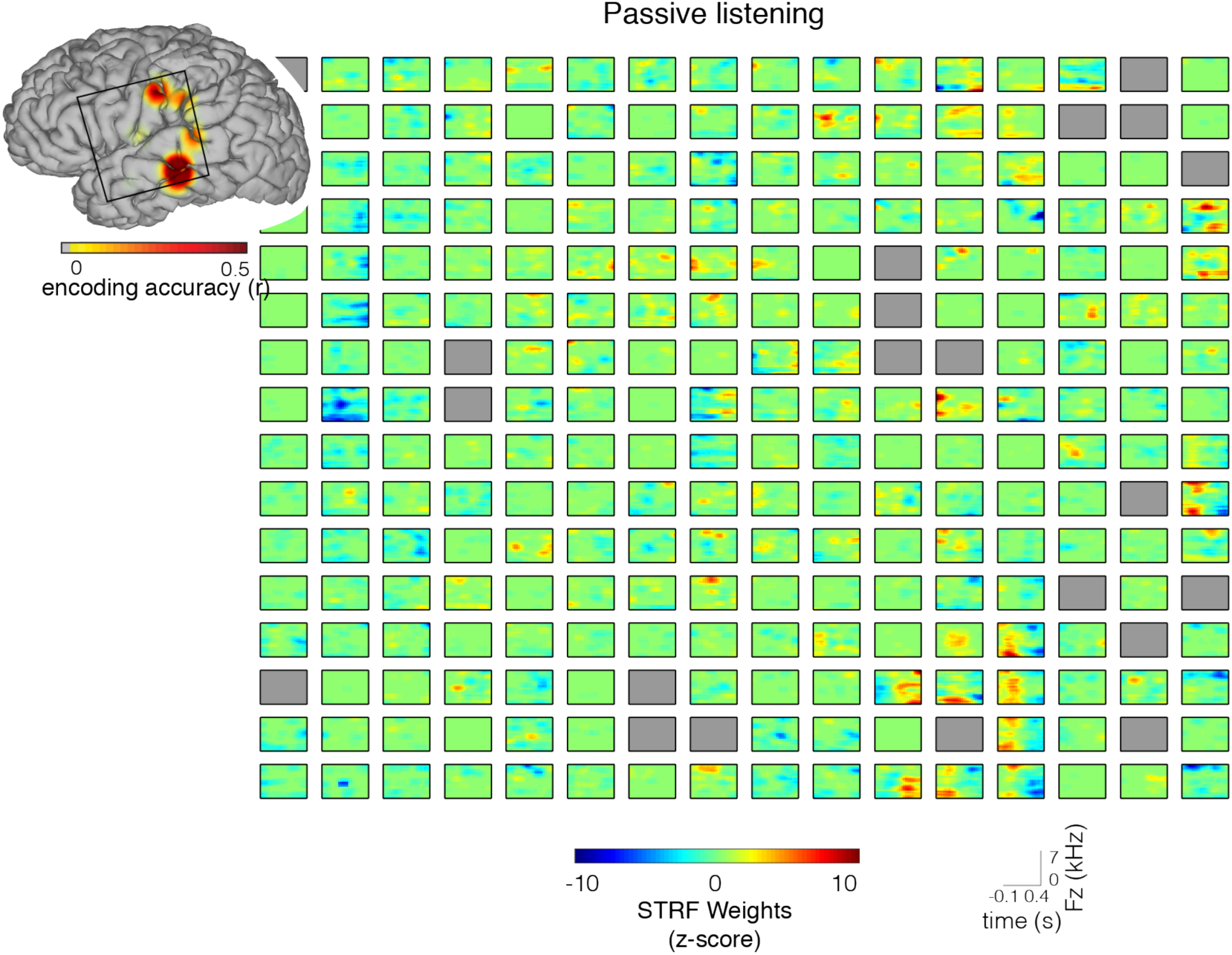
**Neural encoding during passive listening**. Standard STRFs for the passive listening model – built on data recorded while the participant listened passively to speech sentences from the TIMIT corpus [64]. Grey electrodes were removed from the analysis due to excessive noise. Encoding accuracy is plotted on the cortical surface reconstruction of the participant’s cerebrum (map thresholded at p<0.05; FDR correction).

